# Electrophysiological characterisation of iPSC-derived human β-like cells and an *SLC30A8* disease model

**DOI:** 10.1101/2023.10.17.561014

**Authors:** Manon Jaffredo, Nicole A. J. Krentz, Benoite Champon, Claire E. Duff, Sameena Nawaz, Nicola Beer, Christian Honore, Anne Clark, Patrik Rorsman, Jochen Lang, Anna L. Gloyn, Matthieu Raoux, Benoit Hastoy

## Abstract

iPSC-derived human β-like cells (BLC) hold promise for both therapy and disease modelling, but their generation remains challenging and their functional analyses beyond transcriptomic and morphological assessments remain limited. Here, we validate an approach using multicellular and single cell electrophysiological tools to evaluate BLCs functions. The Multi-Electrode Arrays (MEAs) measuring the extracellular electrical activity revealed that BLCs are electrically coupled, produce slow potential (SP) signals like primary β-cells that are closely linked to insulin secretion. We also used high-resolution single-cell patch-clamp measurements to capture the exocytotic properties, and characterize voltage-gated sodium and calcium currents. These were comparable to those in primary β and EndoC-βH1 cells. The K_ATP_ channel conductance is greater than in human primary β cells which may account for the limited glucose responsiveness observed with MEA. We used MEAs to study the impact of the type 2 diabetes protective *SLC30A8* allele (p.Lys34Serfs*50) and found that BLCs with this allele have stronger electrical coupling. Our data suggest that with an adapted approach BLCs from pioneer protocol can be used to evaluate the functional impact of genetic variants on β-cell function and coupling.

**Article highlights:** *Why did we undertake this study?:* iPSC-derived beta like cells (BLCs) from pioneering protocols are known for variable β-cell functionality and mixed cell populations which greatly limits downstream functional assessment. To overcome this challenge, we used electrophysiological tools to provide a detailed functional assessment of BLCs. We then wanted to apply this approach to identify additional functional differences from BLCs carrying a protective Type 2 Diabetes *SLC30A8* allele.

*What is the specific question(s) we wanted to answer?:* Can an electrophysiological approach provide detailed functional characterisation of iPSC-derived BLCs? Is this approach sensitive enough to capture functional differences resulting from *SLC30A8* loss of function (lof)?

*What did we find?:* We found that BLCs generated from pioneer protocol shared electrophysiological features with human pancreatic β-cells, and that a T2D-protective *SLC30A8* lof allele improves the electrical coupling activity of human β-cells.

*What are the implications of our findings?:* Our findings validate the use of intra- and extra-cellular electrophysiology to assess and monitor the functions of BLCs. Our approach opens the perspective of using MEAs to live-monitor the differentiation quality of iPSC-derived BLCs and to determine the functional consequences of diabetes-associated variants.

## Introduction

The differentiation of human inducible Pluripotent Stem Cells (hiPSC) into β-like cells (BLCs) has become the favoured human cellular system to model the effects of specific alleles on beta cell development and function (1–3). However, the difficulties in producing homogenous cultures of matured BLCs have rendered functional characterisation challenging. Live recordings of the extracellular electrical activity of organoids like pancreatic islets on-chip provide an opportunity to address these challenges since β-cell electrical activity is tightly linked to insulin secretion (4, 5). Nutrients stimulate insulin secretion by inhibiting plasmalemmal ATP-regulated K^+^ (K_ATP_) channels leading to membrane depolarization and action potential firing via regenerative activation of voltage-gated Na^+^, Ca^2+^ and K^+^ channels. The electrical activity not only regulates insulin secretion but also influences mitochondrial metabolism and gene expression (4, 6, 7). Hence, monitoring the electrical activity provides rapid and precise functional readout of BLC properties with unequalled high temporal resolution for kinetics analysis.

We used both intra- and extra-cellular electrophysiology to (i) provide a functional assessment of BLCs, and (ii) to study the effect of T2D-risk associated alleles on the function of BLCs. BLCs were generated from a standard protocol of differentiation adapted from Rezania et *al.* (3). This protocol provides BLC preparations with heterogenous cellular population that limits its overall usefulness, making of the BLCs a complex cellular model to use.

Traditional intracellular electrophysiological techniques (i.e. patch clamp) provide detailed information on specific channel activities but is low throughput and technically demanding. Extracellular Multi-Electrode Array (MEA) recordings permit a non-invasive and easily accessible long-term monitoring of the electrical activity of cell clusters (5, 8–10). The MEA captures slow potentials (SPs) (11), an electrical signal reflecting the propagation of electrical activity across a syncytium of islet β-cells. These SPs result from the electrical coupling between β-cells and their frequency correlates with insulin secretion from human isolated pancreatic islets (5, 11).

We examined the functions of BLCs generated by the standard and most commonly used differentiation protocol(3) with a view to exploring their electrophysiological characteristics. We then applied this approach to a proof-of-concept study using BLCs expressing a T2D- protective alleles of the zinc transporter gene *SLC30A8*. Ultimately, our approach is adaptable to BLCs derived by any differentiation protocol and will facilitate the characterisation of alleles associated with or causal for diabetes on β-cell function (12–14).

## Research Design and Methods

### EndoC-βH1 and iPSC-derived β-like cells

Endocell provided the EndoC-βH1 cell line. Cells were regularly tested for mycoplasma contamination (Lonza, LT07-118) and cultured as previously published (15). The parental SB Ad3.1 hiPSC line was previously generated (16) and was subjected to the following quality control checks: SNP-array testing via Human CytoSNP-12 v.2.1 beadchip (Illumina, catalog no. WG-320-2101), DAPI-stained metaphase counting and mFISH, flow cytometry for pluripotency markers (BD Biosciences, 560589 and 560126). CRISPR-Cas9 genome editing was used to generate the *SLC30A8*-p.Lys34Serfs*50 SB Ad3.1 hiPSC line and the resulting cell lines were differentiated using the Rezania protocol (3) as previously described (12).

### Expression assays

Briefly, RNA was extracted using TRIzol Reagent (Life Technologies, 15596026) according to the manufacturer’s instructions. Complementary DNA was amplified using the GoScript Reverse Transcription Kit (Promega, A5000). qPCR was performed using 40 ng of cDNA, TaqMan Gene Expression Master Mix (Applied Biosystems, 4369017) and primer/probes for *SLC30A8* (Hs00545182_m1), or the housekeeping gene TBP (Hs00427620_m1). Gene expression was determined using the ΔΔCT method by normalizing to TBP as previously published (12).

### Secretion assays

iPSC-derived BLC clusters at stage 7 were incubated for 1 hr with glucose-free Krebs-Ringer buffer (KRB) medium consisting of (mM) 138 NaCl, 3.6 KCl, 0.5 MgSO_4_, 0.5 NaH_2_PO_4_, 5 NaHC0_3_, 1.5 CaCl_2_ and 5 HEPES (adjusted to pH 7.4 with NaOH) and supplemented with 0.2% w/v BSA buffer. Depending on the quantity available, preparations were either sequentially stimulated with indicated agents or stimulated in parallel for 20 minutes. Supernatants were taken for determination of insulin release (ELISA, Alpha Laboratories) and the clusters were further processed for electron microscopy.

### Immunocytochemistry

BLCs were fixed in 4% paraformaldehyde, encapsulated in histogel (Thermo Fisher) and embedded in paraffin wax. Dewaxed 5-micron thick sections were blocked with swine serum. Insulin, glucagon and somatostatin were labelled overnight at 4°C using respectively anti-insulin (epitope: human B-chain, in house, 1/500; guinea pig), mouse anti-glucagon (Sigma, 1/500) and rabbit anti-somatostatin (Dako,1/200). The secondary antibodies (anti-guinea-pig 633, anti-mouse TRITC, and anti-rabbit 488) were diluted 1/100.

### Electron microscopy

Each BLC specimen was fixed in 2.5% glutaraldehyde, and either post-fixed in 2% uranyl acetate, dehydrated in graded methanol, and embedded in London Resin Gold (Agar Scientific, Stansted, UK) or post-fixed in 1% osmium tetroxide plus 1.5% potassium ferricyanide in cacodylate buffer, and embedded in Spurr’s resin. Ultrathin LRG sections (70 nm) cut onto nickel grids were immunolabelled with anti-somatostatin (Santa Cruz Biotechnology, #25262, 1:10) followed by anti-rabbit biotin (Vector Laboratories, Peterborough) and streptavidin gold 15 nm (British Biocell International, Cardiff, UK). Insulin was immunolabelled (DAKO, Ely, UK, 1:500) followed by anti-guinea pig gold 10 nm (British Biocell International). Sections were viewed on a Joel 1010 microscope (accelerating voltage 80[kV) with a digital camera (Gatan, Abingdon, UK).

### Intracellular electrophysiology

BLC clusters were dispersed by 3 min trypsin digestion, plated in 35 mm dishes and cultured overnight. Measurements were performed at 32°C in standard whole cell configuration using an EPC-10 amplifier and Pulse software. Exocytosis was measured using membrane capacitance measurements. The extracellular medium was composed of (in mM): 118 NaCl, 5.6 KCl, 2.6 CaCl_2_, 1.2 MgCl_2_, 5 HEPES, and 20 tetraethylammonium (TEA) (pH 7.4 with NaOH). The intracellular medium contained (in mM): 129 CsOH, 125 Glutamic acid, 20 CsCl, 15 NaCl, 1 MgCl_2_, 0.05 EGTA, 3 ATP, 0.1 cAMP, 5 HEPES (pH7.2 with CsOH). Cell size was estimated from the membrane capacitance. Calcium currents were measured from −40 mV to +40 mV and triggered by a 50 ms depolarisation from the resting potential (−70 mV). The mean current 10-50 ms after the onset of the depolarization was used to determine the current amplitude; the initial 10 ms ignored to minimise contribution by rapidly inactivating voltage-gated sodium currents. The membrane potential recordings were obtained using perforated patch. The extracellular medium was composed of (in mM) 138 NaCl, 3.6 KCl, 0.5 MgSO_4_, 0.5 NaH_2_PO_4_, 5 NaHC0_3_, 1.5 CaCl_2_ and 5 HEPES (pH 7.4 with NaOH). The extracellular medium was supplemented with glucose (1 or 20 mM) or tolbutamide (0.2[mM) as indicated. The intrapipette solution contained (in mM) 128 K-gluconate, 10 KCl, 10 NaCl, 1 MgCl_2_ and 10 HEPES (pH 7.35 adjusted with KOH). Perforation of the membrane was achieved using amphotericin B (0.24[mg/ml)(6).

### Extracellular electrophysiology

Recordings with MEAs were performed as published(5) at 37° C either in a solution containing (in mM): NaCl 135, KCl 4.8, MgCl_2_ 1.2, CaCl_2_ 2.5, HEPES 10 and glucose as indicated (pH 7.4 adjusted with NaOH) or in a more complete medium (MCDB131) without cytokines. Membranes (ALA-MEA-MEM-PL, MCS) were used to cover the MEA surface to suppress evaporation as published (11). Pictures of BLCs on MEAs were taken before and after each experiment in order to localize electrodes covered with cells. Extracellular field potentials were acquired at 10 kHz per electrode, amplified and filtered (analog) at 0.1-3000 Hz with a USB-MEA60-Inv-System-E amplifier (Multichannel Systems; gain: 1200) controlled by MC_Rack software (Multichannel Systems). Data were analysed either offline or online. For offline analysis, MC_Rack software was used to isolate SPs using a 0.2–2 Hz band-pass digital filter (Butterworth 2^nd^ order). For determination of frequencies, SPs were detected using the threshold module of MC_Rack with a dead time (minimal period between two events) set to 300 ms. On-line hard real-time acquisition and data processing were performed with our configurable acquisition board as published (9).

## Analysis and software

Data are presented as means values and SEM. The number of experiments and details of the statistical analysis are in the figure legends. For electrophysiology and microscopy, n represent the number of cells from several preparation or passages of the same batch. For insulin secretion assay, n represents the independent experiments from different batches of differentiation. For MEA data, n represents the number of electrodes and N the number of preparations. Data were analysed using R software, OriginPro 2020, and GraphPad Prism (V8.0.1).

## Results

### Generation of hiPSC-derived BLCs

The BLCs used for the functional characterization were obtained following the 25-day differentiation protocol adapted from Rezania *et al.* (3). (Supplementary Figure 1A). BLC were either generated in 2D (monolayers) or in 3D (air-liquid interface) as clusters. Both 2D and 3D cultures expressed key genes involved in β-cell maturation (*MAFA* and *PDX1)*, pancreatic hormones (*INS, GCG, SST*) and β-cell function such as K_ATP_ channel subunits (*ABCC8* and *KCNJ11*) (Supplementary Figure 1B).

We stained preparations for pancreatic hormones to ascertain the differentiation efficiency (Supplementary Figure 1C). As previously described (3), cells were positive for pancreatic endocrine hormones with the majority positive for insulin, while some BLCs were polyhormonal: positive for glucagon or somatostatin in addition to insulin (Supplemental Figure 1C, dashed circles). These observations were confirmed by immunogold labelling of insulin (white arrowheads) and glucagon (black arrowheads) which showed distinct α-like and β-like cells with immunogold particles in vesicular structures (Supplemental Figure 1D).

Some cells presented multiple types of endocrine-like vesicles such as typical insulin-positive and glucagon-positive vesicles, the latter composed of a limited halo and a two-phased electron-dense core (Supplemental Figure 1E, white arrowheads: insulin-like vesicles, black arrowhead: glucagon-like vesicles). Insulin- and somatostatin-containing polyhormonal cells were also present with the two hormones in distinct vesicles as well as within the same vesicles (Supplemental Figure 1F, white star). We also detected typical insulin containing vesicles presented an electron-dense central core surrounded by a clear halo (Supplemental Figure 1G, white arrows). Large dense core vesicles were 200 to 500 nm in diameter with an average of 290 nm for monolayers and of 310 nm for clusters (Supplemental Figure 1H).

Consistent ultrastructure observations, BLCs expressed marker genes of vesicular trafficking such as chromogranin A (*CHGA*), markers of the maturation of insulin such as the proprotein convertase subtilisin/kexin type 1 (*PCSK1*) (Supplemental Figure 1I). Overall, the differentiation efficiency observed here is in line with previously reported outcomes for this(3) and other protocols (17–21).

### Multicellular activity measured with MEAs

In islets, β-cells are electrically coupled via gap junctions and function as a syncytium(5, 8, 9, 11, 22). The electrical coupling properties can be monitored non-invasively by culturing BLCs directly on the electrodes of MEAs (Figure 1A). As human islet β-cells, the BLCs exhibited action potential (AP), seen as discrete short-lived spikes, and slow potentials (SPs) (Figure 1B). SPs are β-cell specific electrophysiological markers and result of the summation signals due to cell-cell coupling via connexin 36 (5, 11). Most BLC preparations (76.5%) were electrically active and generated both SPs and APs (Figure 1C) and differentiation in clusters vs. in monolayers yielded similar results. However, SPs were more frequently observed in monolayers (Supplementary Figure 2A). This was consistent with the greater expression of Gap Junction protein Delta 2 (*GJD2)* (Supplemental Figure 2B) which is the dominant connexin expressed in human β- cells and is essential to the propagation of the SP signals.

**Figure 1:**
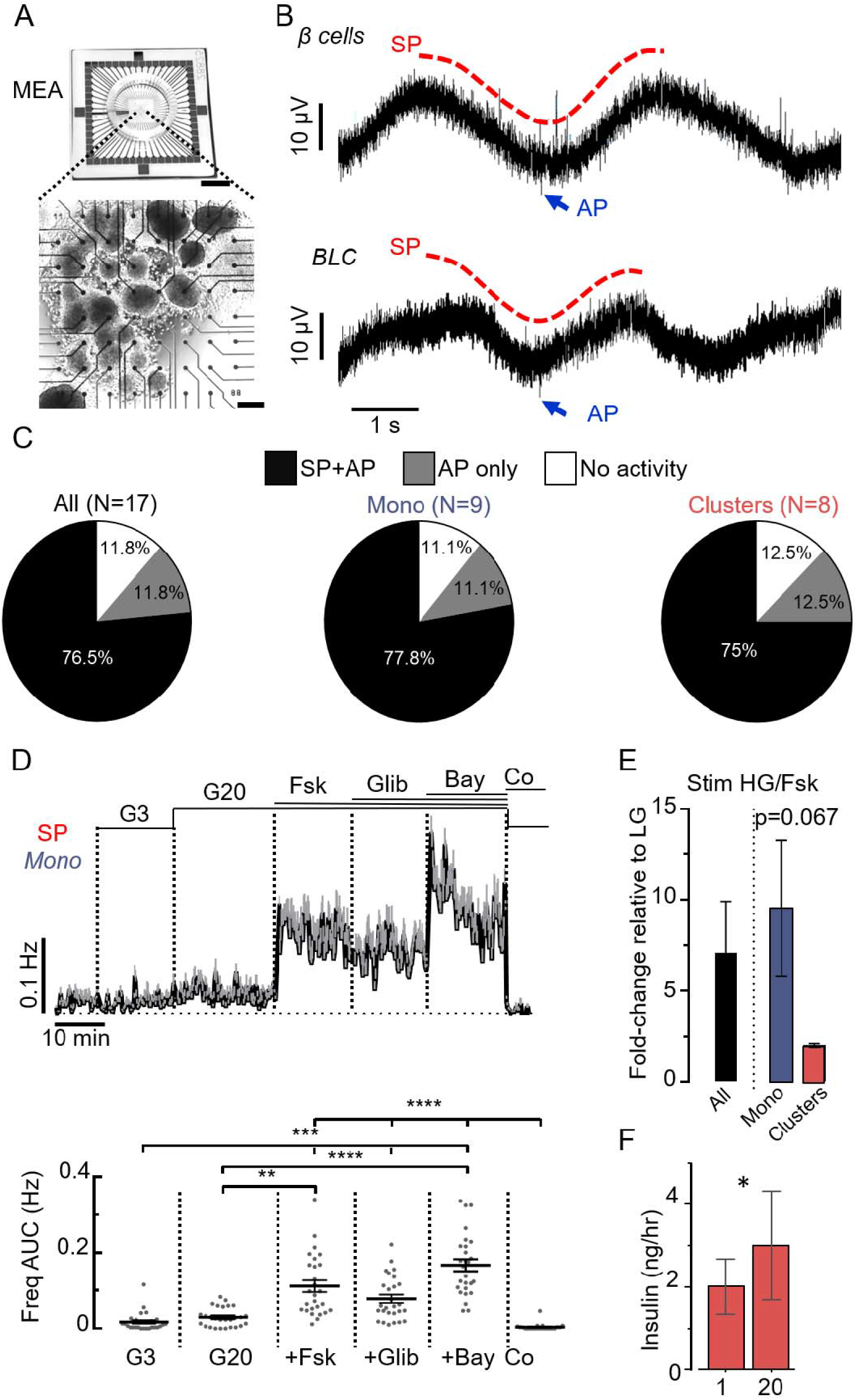
Multicellular MEA analysis of BLCs reveals electrical coupling. **A.** Picture of MEA (top, scale bar: 1 cm) with BLCs (in clusters) cultured on the 59 electrodes (bottom, scale bar: 200 µm). **B**. Representative recordings of the electrical activity of primary human islet β cells (top) and BLCs (bottom) obtained from one of the MEA-electrodes. The two characteristic types of electrical signals were present in both cells: the multicellular SPs (examples in red) and the unicellular APs (examples in blue). See also Supplementary Fig. 2 for comparison with recordings of EndoC-βH1 cells. **C.** Inter-preparation analysis. Proportion BLC preparations presenting SPs and APs (black), only APs (grey) or neither SPs nor APs (white). Data for all preparations (All: clusters and monolayers), monolayers only (Mono) and clusters only are presented. **D.** MEA-based functional quality control of a preparation of BLCs in monolayer (Mono). Variations of SP frequencies (top: means values ±SEM; bottom: statistics performed on areas under the curves (AUC) normalized over time; last 5 min for G3 and first 5 min for the other conditions) in response to several stimuli: 3 mM glucose (G3), 20 mM glucose (G20) alone, followed by successive additions of forskolin (Fsk, 1 µM), glibenclamide (Glib, 100 nM) and Bay K8644 (Bay, 10 µM). At the end of the protocol, G3 with CoCl_2_ (Co, 2 mM) was applied to inhibit the electrical activity. (n=27; **p<0.01, *** p<0.001, **** p<0.0001; Friedman test). See also Supplementary Fig. 3 for the same quality control on a preparation of BLCs in clusters. **E.** Electrical responses to high glucose and Fsk (HG/Fsk) were represented as fold change of AUC SP frequency relative to low glucose (LG), and were higher in monolayers than in clusters (Mono, N=4; Clusters, N=2; Welch’s t-test). **F.** Glucose stimulation (20mM) in clusters preparations elicited a mild but significant increase in insulin secretion (N=5, 2p=0.025, two-tail ratio paired t-test).

Notable differences in SPs were observed in response to β cells activators. In monolayers SPs were rare at 3mM glucose with transient and discrete increase in their frequency at 20 mM glucose (Figure 1D), and their frequency significantly increased by 7-fold in the presence of the adenylate cyclase activator forskolin. The effect of forskolin was more pronounced in monolayers than in clusters (Figure 1D and E, Supplemental Figure 3). The K_ATP_ channel blocker glibenclamide did not increase SP frequency beyond that produced by forskolin in both monolayers (Figure 3D, Supplemental Figure 3). However, a further increase was seen in the presence of the L-type Ca^2+^ channel activator Bay K8644. Inhibition of Ca^2+^ channels by Co^2+^ abolished the SPs (Figure 1D, Supplemental Figure 3), confirming their dependence on voltage-gated Ca^2+^ channels as previously reported in islets(11). High glucose concentration (20 mM) only increased insulin secretion by 50% in BLC clusters (Figure 1F) echoing the weak effect on electrical activity (Figure 1D). Monitoring BLC in monolayers using an automatic real time analysis of SP frequency (8, 9) revealed that similar (50%) Glucose-Stimulated Insulin Secretion (GSIS) was associated with a dramatic but transient stimulation of the SP frequency (analysed as in (8, 9)). These differences in SP frequency are in line with *GJD2* in both BLCs models (Supplementary Figure 2).

### Single-cell electrophysiological properties of BLCs

To further detail the electrophysiological properties of BLCs, we monitored their membrane potential using perforated patch whole-cell measurements. In a minority of cells, increasing glucose from 1 to 20 mM induced membrane depolarization and increased AP firing (Figure 2A) as expected from fully functional β-cells. The APs were triggered at >-40mV (Figure 2A, red inset) and their shapes and duration mirrored those reported for primary human β cells and the human beta cell line EndoC-βH1 cells(6, 23, 24). For the majority of the cells however, while the membrane potential oscillated from -60 to -40mV, neither glucose nor tolbutamide promoted AP firing (Figure 2B red inset). In these cells, the average conductance in 1 mM glucose was variable and averaged to 215.7±80.5 pS.pF^-1^ (Figure 2C). This is considerably larger than the conductance previously reported for human primary β-cells and EndoC-βH1 (∼60pS.pF^-1^) (6, 23, 24). When normalised to the initial conductance measured with 1 mM glucose, the stimulation with either 20 mM glucose and with addition of 0.2 mM tolbutamide reduced the K_ATP_ conductance by 50% and 80% respectively (Figure 2C, right).

**Figure 2:**
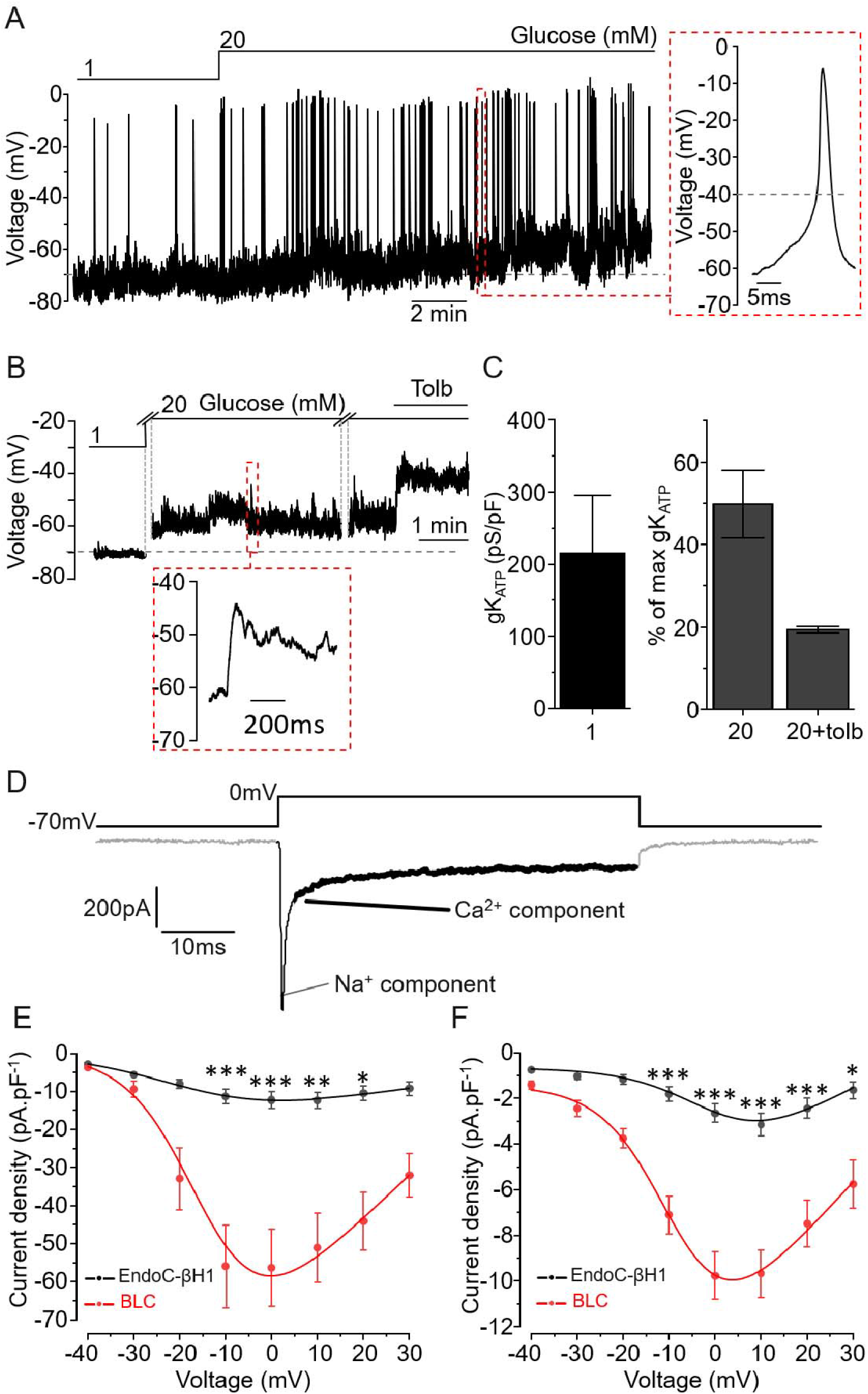
Single-cell electrical properties of BLCs. **A.** BLC membrane potential recording in an active cell upon incubation with 1 and 20mM glucose and representative action potential (red inset). **B.** Representative recording of a silent cell. Increase glucose concentration promoted slight membrane depolarisation but did not translate in AP firing (red inset) **C (left).** K_ATP_ channel conductance of silent BLCs at 1mM glucose. **C (Right).** K_ATP_ channel conductance was reduced by 50% during incubation with 20 mM glucose and further reduced by 80% when 0.2 mM tolbutamide was added (n=3 cells in C). Data are normalised to the initial conductance measured with 1 mM glucose. **D**. Representative trace of current elicited by a square depolarisation from the resting potential (-70 mV) to 0 mV. The inward current is composed of a rapid (<5 ms) sodium component (thin black section) followed by a sustained calcium component (black section in bold). **E-F.** Current-voltage relationship for the sodium (E) and calcium (F) components. Black: EndoC-βH1 (n=10 cells) and BLC in red (n=17 cells).* p<0.05, ** p<0.01, *** p<0.001 Paired comparison and Tukey.

We measured the voltage-activated currents that underlie APs and trigger exocytosis (Figure 2D). Like in primary β-cells (23), the current evoked by 50 ms depolarisation contained (i) a rapidly inactivating component (reflecting the activation of voltage-gated Na^+^ channels) and (ii) a sustained component reflecting the activity of voltage-gated Ca^2+^ channels. We compared their current properties to the well-established human β-cell model EndoC-βH1 (6, 15). In both cell types, Na^+^ (Figure 2E) and Ca^2+^ (Figure 2F) current densities peaked during depolarizations to 0 mV were significantly greater in BLC than in EndoC-βH1 and close to values reported in primary human β-cells(23).

### Regulated exocytosis in BLCs

To further characterise the secretory capacities of BLCs, we used high-resolution membrane capacitance measurements (25, 26). This approach measures the surface area of the cells which increases in proportion to the number of vesicles releasing their content through regulated exocytosis. To estimate the kinetics of exocytosis, the cells were first subjected to depolarisations (from -70 to 0 mV) with duration ranging from 10 to 800 ms (Figure 3A). BLCs and EndoC-βH1 presented similar kinetics, reaching a maximum amplitude of exocytosis of ∼10 fF.pF^-1^ (Figure 3B, inset). For the shortest depolarisations (<50 ms), the amount of depolarising charges (such as Ca^2+^ entry) was compared with the level of exocytosis triggered (Figure 3C). In both cell-types the amplitude of exocytosis strongly correlated with Ca^2+^ entry. However, the correlation was significantly right-shifted in BLCs. Thus, Ca^2+^ entry appears more tightly coupled to exocytosis in EndoC-βH1 cells.

**Figure 3:**
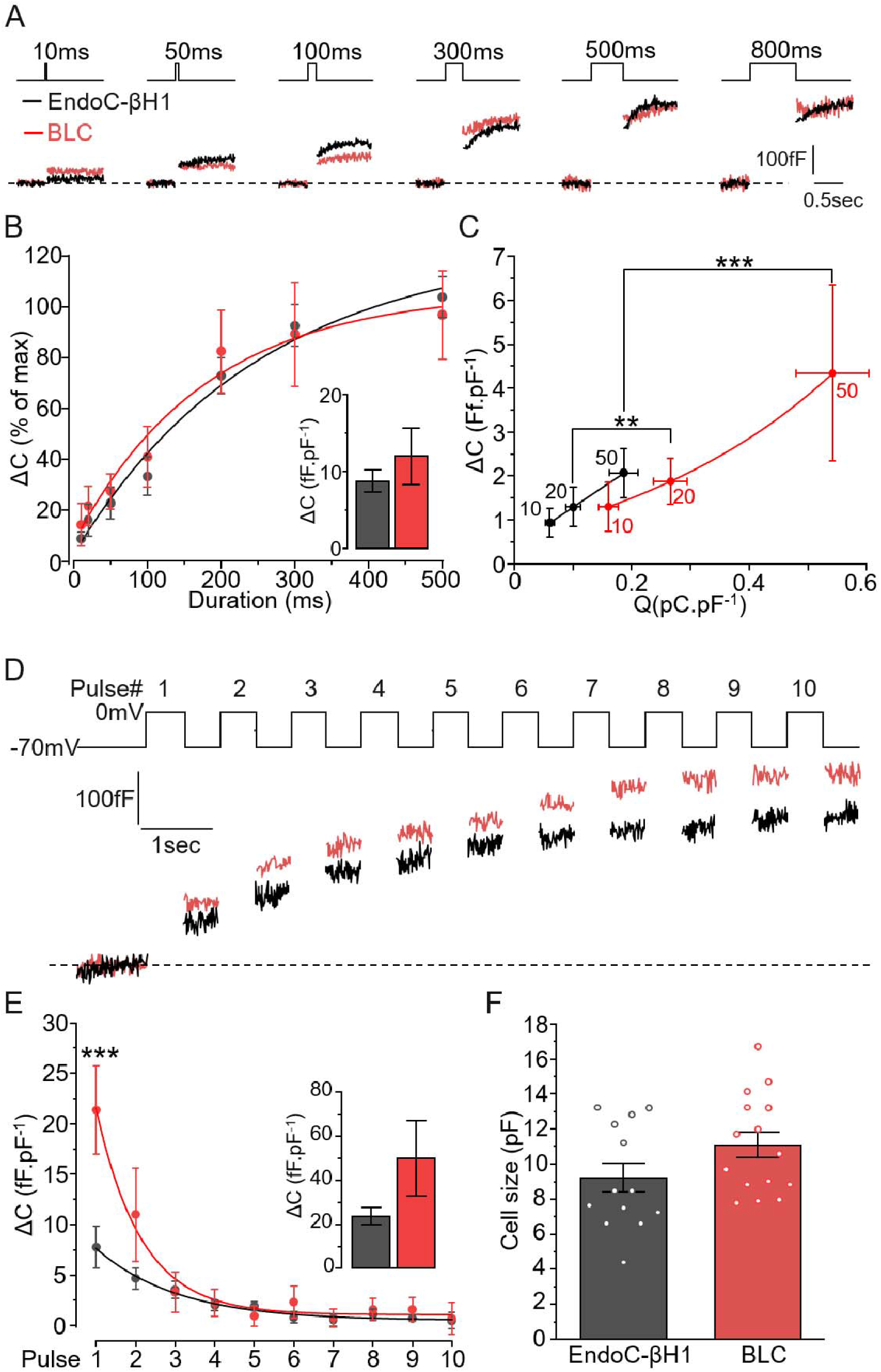
Comparison of the exocytotic properties of BLCs and EndoC-βH1 cells. **A-B.** EndoC-βH1 (black) and BLCs dispersed from cluster preparations (red) were subjected to increased durations of depolarisation (from 10 ms to 800 ms) and the resulting exocytosis events were measured as an increase in cell surface area. **B.** The stimulations triggered similar kinetics of exocytosis in both models with a plateau reached from 300ms depolarisation. Inset: maximum exocytosis elicited by 800ms stimulation (n=13 and n=15 cells for EndoC-βh1, black, and BLC, red, respectively). **C.** Charge-exocytosis relationship in EndoC-βH1 and in BLC. The right-shifted correlation in BLCs is driven significantly by the amount charges. **D.** Representative traces of the maximum exocytosis elicited by ten depolarisations of 500 ms. **F.** Quantification of exocytosis increment elicited at each pulse (n=13 and 11 cells for EndoC-βH1 and BLC, respectively). Insert: total cumulative exocytosis measured at pulse 10. **G.** The initial size of the cells (pF) were similar between the two β-cell models (n=13 and n=15 cells for EndoC-βh1, black, and BLC, red, respectively). Statistics, **p<0.01, *** p<0.001, Paired comparison and Bonferroni posthoc test.

We then submitted BLCs to a train of 10 depolarisations of 500 ms at 1 Hz to trigger the maximal exocytotic response (representative traces, Figure 3D). The increase in membrane capacitance per pulse was biphasic with the largest response observed during the initial 3 pulses (Figure 3E). The initial stimulation (during the first pulse) promoted significantly greater exocytotic response in BLCs. Although consistently of greater amplitude in BLCs, the total exocytosis elicited at the end of the 10 depolarising pulses remained statistically similar in both cell types (Figure 3E, inset). Finally, the differences in current densities (Figure 2) as well as in exocytosis (Figure 3) between BLCs and EndoC-βH1 were independent of variations in cell size (Figure 3F).

### A diabetes protective allele improves BLCs’ electrical responses

We then used the MEA to examine the effect of a T2D-associated allele on BLCs function. As a proof-of-concept, we measured the electrical behaviour of BLCs derived from a loss of function *SLC30A8* allele (p.Lys34Serfs50*) using CRISPR-Cas9 genome-edited hiPSCs (12). Several alleles of the vesicular zinc transporter ZnT8 (*SCL30A8*) strongly influence the risk of developing T2D (12, 27). The frameshift allele p.Lys34Serfs50* provides protection against T2D and leads to loss of *SLC30A8* expression by Nonsense Mediated Decay (NMD) in homozygous BLCs (12, 27, 28). The electrical activities of BLCs generated in monolayer from the CRISPR-edited *SLC30A8* p.Lys34Serfs50* was compared to the unedited control cells (Sham) (Figure 4). In both cell lines, electrical activity was unaffected by a small increase in glucose (3 to 10 mM) (Figure 4A). Addition of forskolin or glibenclamide induced a biphasic and greater increase of SP frequency in the p.Lys34Serfs50* line but not in control BLCs (Figure 4A, B). Finally, incubating BLCs with 10 mM glucose in complete culture medium (in presence of amino acids and additional nutrients) significantly increased SP frequency in p.Lys34Serfs50* BLCs but was without effect in the CRISPR-sham control line (Figure 4C and D). This suggests that additional metabolic pathways are improved upon *SLC30A8* loss of function in β cells, which is in line with the T2D-protection associated with *SLC30A8* loss of function in humans.

**Figure 4:**
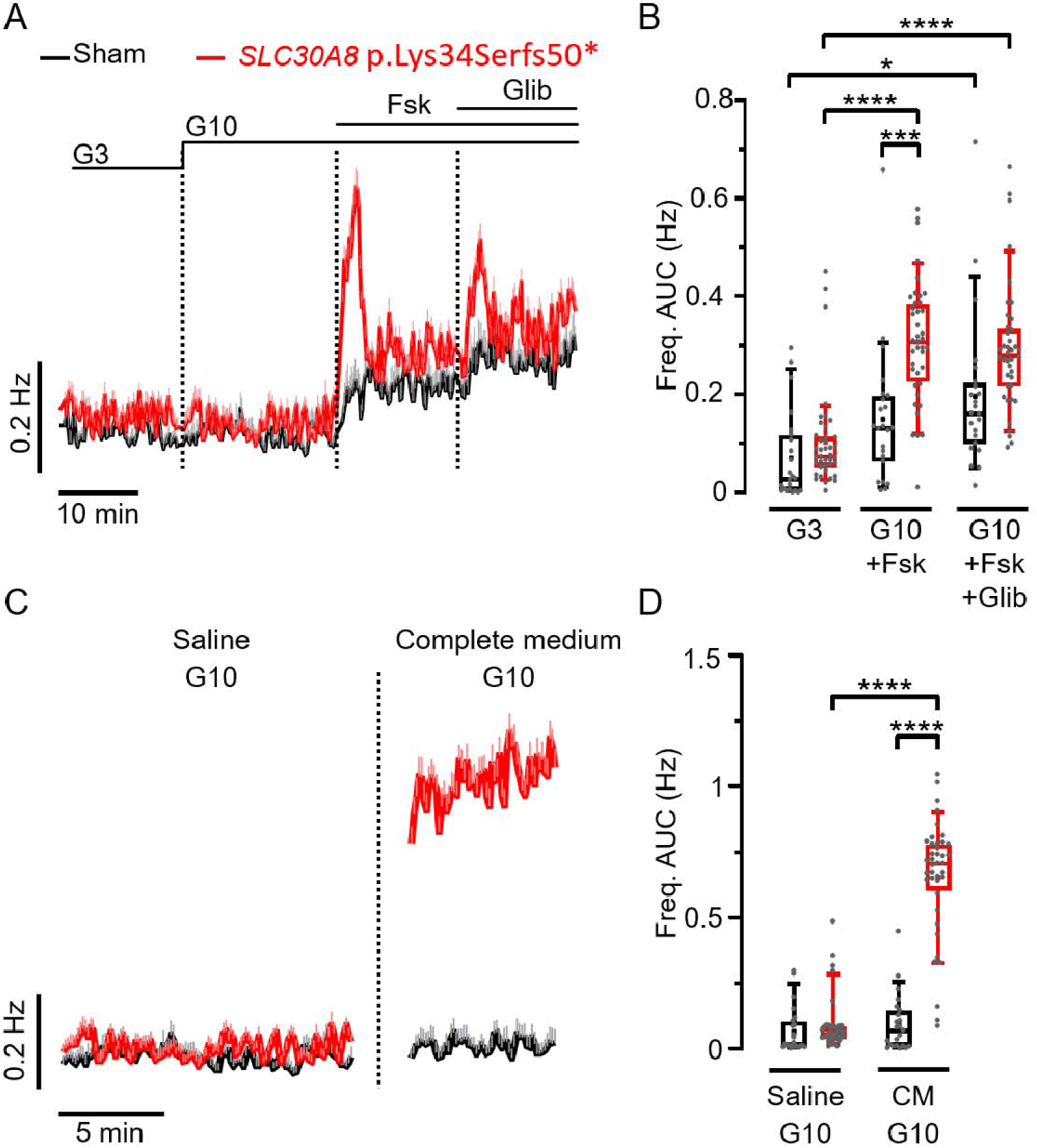
The loss-of-function mutation of *SLC30A8* increases slow potential frequencies in CRISPR-edited BLC monolayers. **A**. Kinetics of average SP frequencies measured in CRISPR-Sham (black) and *SLC30A8*- edited BLCs (red) in the presence of 3 mM (G3), and 10 mM (G10) glucose alone followed by successive additions of forskolin (Fsk, 1 µM) and glibenclamide (Glib, 100 nM, in the presence of Fsk). **B**. Statistics on AUCs of SP frequencies (normalized over time; last 5 min for G3 and first 5 min for the other conditions) obtained from data in (A) (n=23-40; N=2 independent mutated clones; *p<0.05, *** p<0.001, **** p<0.0001; Kruskal-Wallis test). **C**. SP frequencies (means +SEM) measured on the same BLCs in the presence of a saline solution and of complete medium containing both 10 mM glucose (G10). **D**. Statistics on AUCs of SP frequencies (normalized over time) of data in C (n=23-40; N=2; **** p<0.0001; Kruskal-Wallis test).

## Discussion

Recent protocols for hiPSC-derived BLC differentiation have opened new perspectives for *in vitro* human disease modelling as well as autologous diabetes cell replacement therapy (29). While these protocols have improved the functional quality of BLCs, modified pioneer protocols remain widely used (12, 30) and dynamic functional analyses are needed to better characterize and standardize these preparations (31). Our data indicate that the secretory vesicle ultrastructure and the exocytotic process in BLCs resemble those in primary human β- cells (4, 23, 25, 32). The properties of the voltage-gated Na^+^ and Ca^2+^ currents in BLCs and the characteristics of APs (when generated) were likewise comparable to human primary β- cells. These data suggest that in BLCs, the exocytotic machinery required to release insulin is expressed, correctly located and functional (23, 33).

As reported previously by others using a similar protocol (3, 17–21), GSIS was clearly of small amplitude in BLCs. Several observations here may explain the difference between exocytosis evoked by electrical depolarisation versus glucose-induced stimulation. First, K_ATP_ channel activity (expressed as membrane conductance) was considerably higher than that of mature β-cells. Although it was reduced upon glucose stimulation, it remained ∼100 pS.pF^-1^, which is likely sufficient to keep the cells repolarised. Second, the presence of bihormonal cells (positive for insulin and somatostatin) releasing a mixed cargo could exert a paracrine inhibitory effect through the activation of somatostatin receptor (6, 34, 35). These poly-hormonal cells, by retain foetal properties (36, 37)] and thus reducing BLC responsiveness, could contribute to low GSIS (38, 39). Third, extracellular electrophysiology detected SPs that reflect β-cell synchronization and strongly advocate that BLC preparations have the capability of behaving like islet micro-organs rather than the sum of isolated cells (5, 11). However, SPs frequency was low compared to primary β cells (5,8,9,11) suggesting a reduced coupling activity that could consequently promote a higher basal secretion and a reduced GSIS (40).

The loss of function allele p.Lys34Serfs50* in *SLC30A*8 leads to increased glucose responsiveness and insulin secretion in human β-cells (12). Although glucose alone did not promote an increased electrical activity, addition of forskolin or glibenclamide, as well as a stimulation with a more physiological medium (i.e. amino acids containing culture medium) had a more potent effect on *SLC30A*8 KO than on control BLCs. This is in line with recent investigation showing *SLC308* KO improved the maturation of hESC-derived β cells (41). As coupling and SPs correlate to insulin secretion (5), our data provide an electrophysiological explanation for the enhanced GSIS *in-vitro* and is in line with the reported reduction in K_ATP_ conductance associated with *SLC30A8* loss of function (12).

Our data suggest a relative immaturity of BLCs especially in the proximal (K_ATP_ channel- dependent) but not in the more distal steps of the stimulus-secretion coupling. This is in line with previous observations on low mitochondrial activities in BLCs resulting from weak TCA anaplerotic pathways (19, 42). Potentiators of glucose stimulation, such as forskolin (raising cytosolic cAMP), significantly increased the electrical response of BLCs, suggesting that components of the glucose amplification pathway are operational (43).

Our results also suggest that functional measurements as well as gene expression should be used to follow the maturation of BLCs. Although we detected *bona fide* markers of β-cell maturation in BLCs, the better response to sulfonylureas than to glucose echoes foetal β-cells properties (44). This reignites the debate as to whether markers such as *MAFA* levels represent a good marker of human BLC maturity (45) and highlights the challenge of predicting β-cell function and maturity from transcriptomic data. Clearly, the best measure of BLC maturity remains in the capacity to respond to physiological concentrations of glucose and other secretagogues capturing the molecular mechanism reported in human pancreatic islets (membrane depolarisation, coupling, and stimulation of insulin secretion).

In conclusion, our data illustrate that functional assessment of BLCs requires a multifaceted approach combining measurements from bulk, micro-organ and single-cells. Our approach paves the way for functional investigations in BLCs and offers the possibility to be integrated in a drug screening pipeline. The electrophysiological characterisation of BLCs, as done here, provides a starting point to understand functional characteristics of the different maturation stages of BLCs and highlights its usefulness to investigate the cellular consequences of specific alleles in human β cells.

## Acknowledgements

Contribution: M.J., N.A.J.K, C.E.D., N.B., C.H., A.C, B.C., S.N., M.R., B.H.: collected the data. P.R., J.L., A.L., A.C, N.B., C.H., C.E.D., N.A.J.K.: edited the manuscript; M.R, B.H: equal contribution (designed the study, analysed data and wrote the manuscript).

Disclosures: ALG’s spouse is an employee of Genentech and holds stock options in Roche. NLB was an employee of University of Oxford when all work contained in this manuscript was conducted. CH is an employee and stock holder of Novo Nordisk A/S.

Funding: BH is a Diabetes UK R.D. Lawrence Fellow (BDA number:19/0005965). M.J. is supported by an EFSD Albert Renold award, MJ & MR are supported by the French Ministry of Research & Education, JL is supported by the FEDER DIAGLYC & ANR DIABLO (ANR-18-CE17-0005). ALG is a Wellcome Trust Senior Fellow in Basic Biomedical Science. Work in ALG’s laboratory was funded by the Wellcome Trust (200837), and the National Institutes of Health (UM-1DK126185). M.R. & B.H. are the guarantors of this study.

