## Supplementary Figure 1 for "Electrophysiological characterisation of iPSC-derived human β-like cells and an *SLC30A8* disease model"

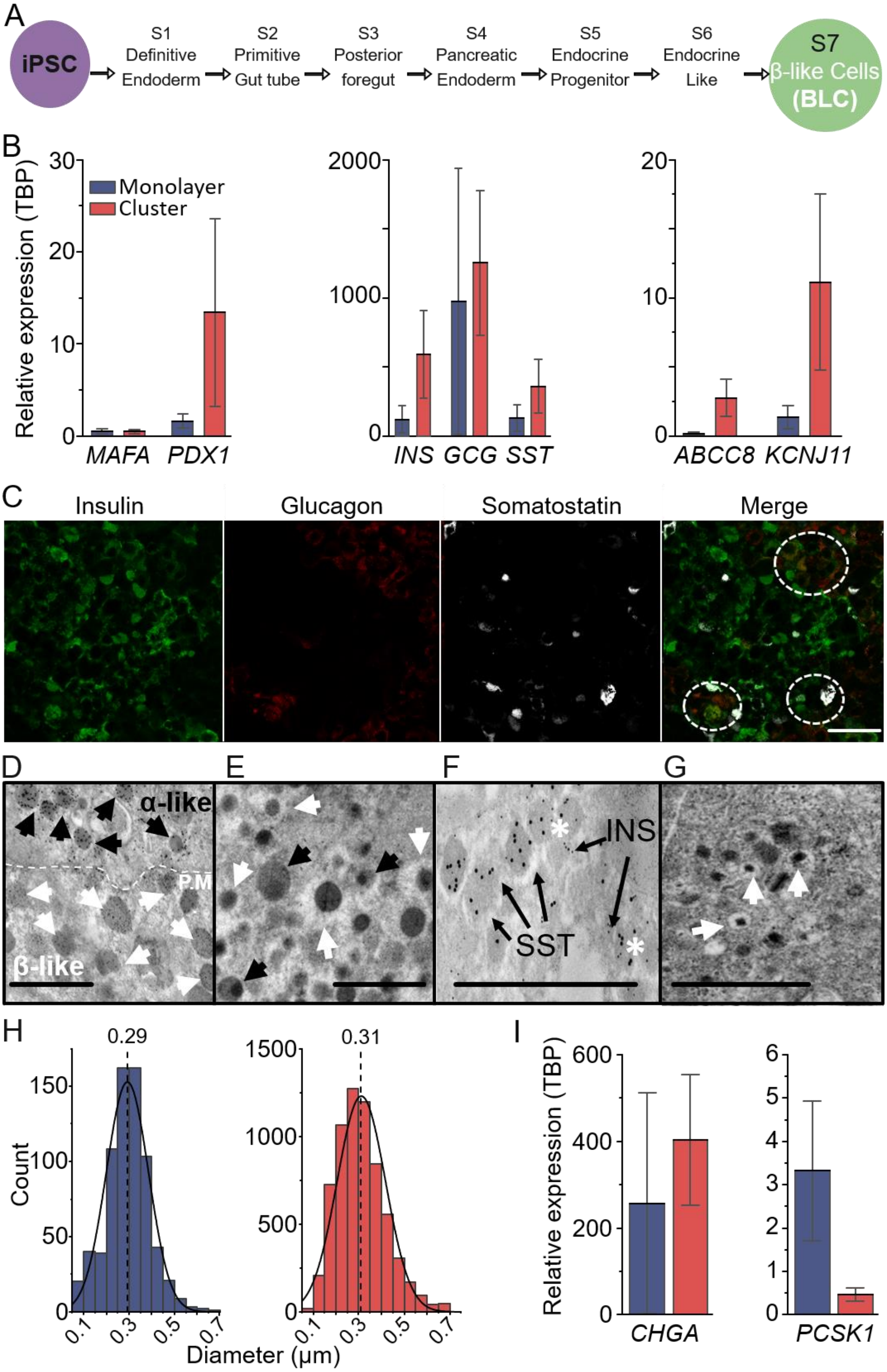


**Supplementary Figure 1**

**Supplementary Figure 1: Overview of the characterisation of iPSC derived BLCs. A.** iPSC were derived following the 25-day differentiation protocol. **B.** RT-qPCR data shows variability in the expressions of differentiation markers (left, $n_{cl}$=8, $n_{mono}$=4), pancreatic hormones (centre, $n_{cl}$=8, $n_{mono}$=4-5), and $K_{ATP}$ channels subunits (right, $n_{cl}$=9, $n_{mono}$=4). **C.** Clusters (n=5) and monolayer (n=1) preparations were stained for pancreatic hormones: Insulin (green), Glucagon (red), and somatostatin (white). Polyhormonal cells are circled (dash circled) on the merge panel. Scale bar 50µm. **D-G**, ultrastructure of BLCs. **D.** Immunogold labelling of cells expressing either insulin (10nm gold particles, white arrowhead, BLC) or either glucagon (15nm gold particles, black arrowheads, α-like cell). PM, Plasma Membrane (dash line). **E.** Electron micrograph of a polyhormonal cell presenting heterogeneous vesicular structures, some typical of glucagon containing vesicles (black arrow heads) or of insulin containing vesicles (white arrowheads). **F.** Immunogold labelling of a polyhormonal cell positive for insulin (10nm gold particles) and somatostatin (SST, 15nm). The hormones could be detected in independent and within the same vesicles (white stars). **G.** Representative electron micrographs of cells containing structurally matured insulin vesicles (white arrows). Scale bar 1µm. **H.** Vesicle size distribution in clusters and monolayers ($n_{mono}$=713 and $n_{cl}$=6532 vesicles). **I.** BLCs express components of the trafficking pathway *CHGA* (left), component of insulin maturation *PCSK1*(right) ($n_{cl}$=6, $n_{mono}$=3).
