## Supplementary Figure 2 for "Electrophysiological characterisation of iPSC-derived human β-like cells and an *SLC30A8* disease model"

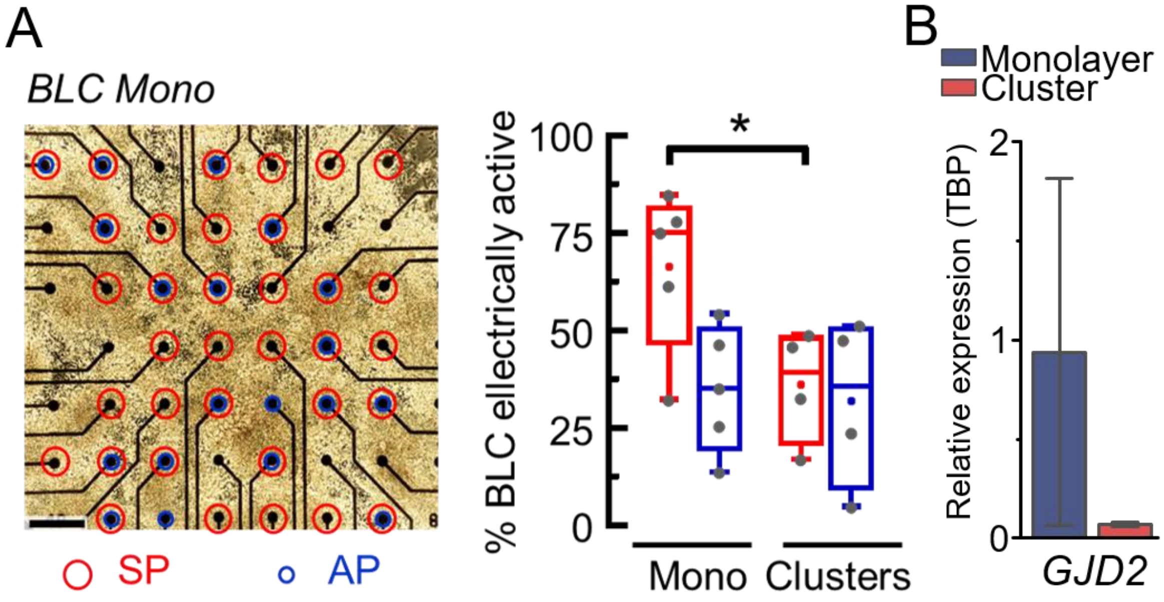


**Supplementary Figure 2**

**Supplementary Figure 2: Intra-preparation analysis of SPs with MEAs and *GJD2* expression in monolayer vs. clusters.** (**A**) Intra-preparation analysis. Left: monolayer of BLCs on a MEA. The electrodes that detected SPs are circled in red and those detecting APs are circled blue. For each experiment, some electrodes could record both or one or none of these electrical. Right: proportion of electrodes with SPs (red) or APs (blue) for each preparation (All, N=9 MEAs; Mono, N=5 MEAs; Clusters, N=4 MEAs; *p<0.05; t-test). (**B**) BLCs express cellular coupling marker *GJD2* essential in the propagation of SPs.
