## Supplementary Figure 3 for "Electrophysiological characterisation of iPSC-derived human β-like cells and an *SLC30A8* disease model"

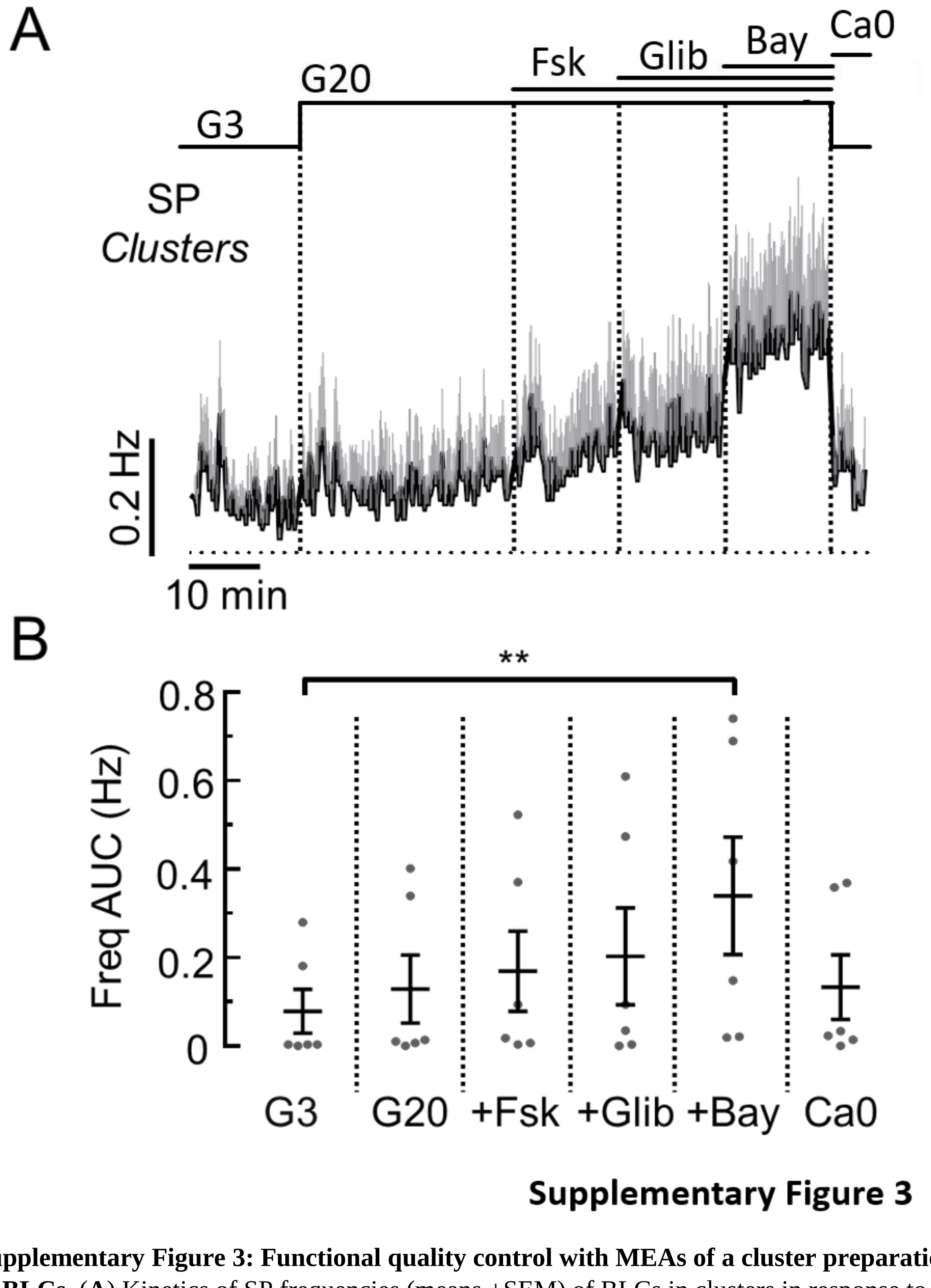


**Supplementary Figure 3: Functional quality control with MEAs of a cluster preparation of BLCs.** (**A**) Kinetics of SP frequencies (means +SEM) of BLCs in clusters in response to several stimuli: 3 mM glucose (G3), 20 mM glucose (G20) alone, followed by successive additions of forskolin (Fsk, 1 µM), glibenclamide (Glib, 100 nM) and Bay K8644 (Bay, 10 µM). At the end of the protocol, G3 without calcium (Ca0) was applied to inhibit calcium entry (n=10). (**B**) Statistics on AUCs of SP frequencies (normalized over time) measured in (**A**) (n=10, **p<0.01; Friedman test).
