## Supplementary Figure 4 for "Electrophysiological characterisation of iPSC-derived human β-like cells and an *SLC30A8* disease model"

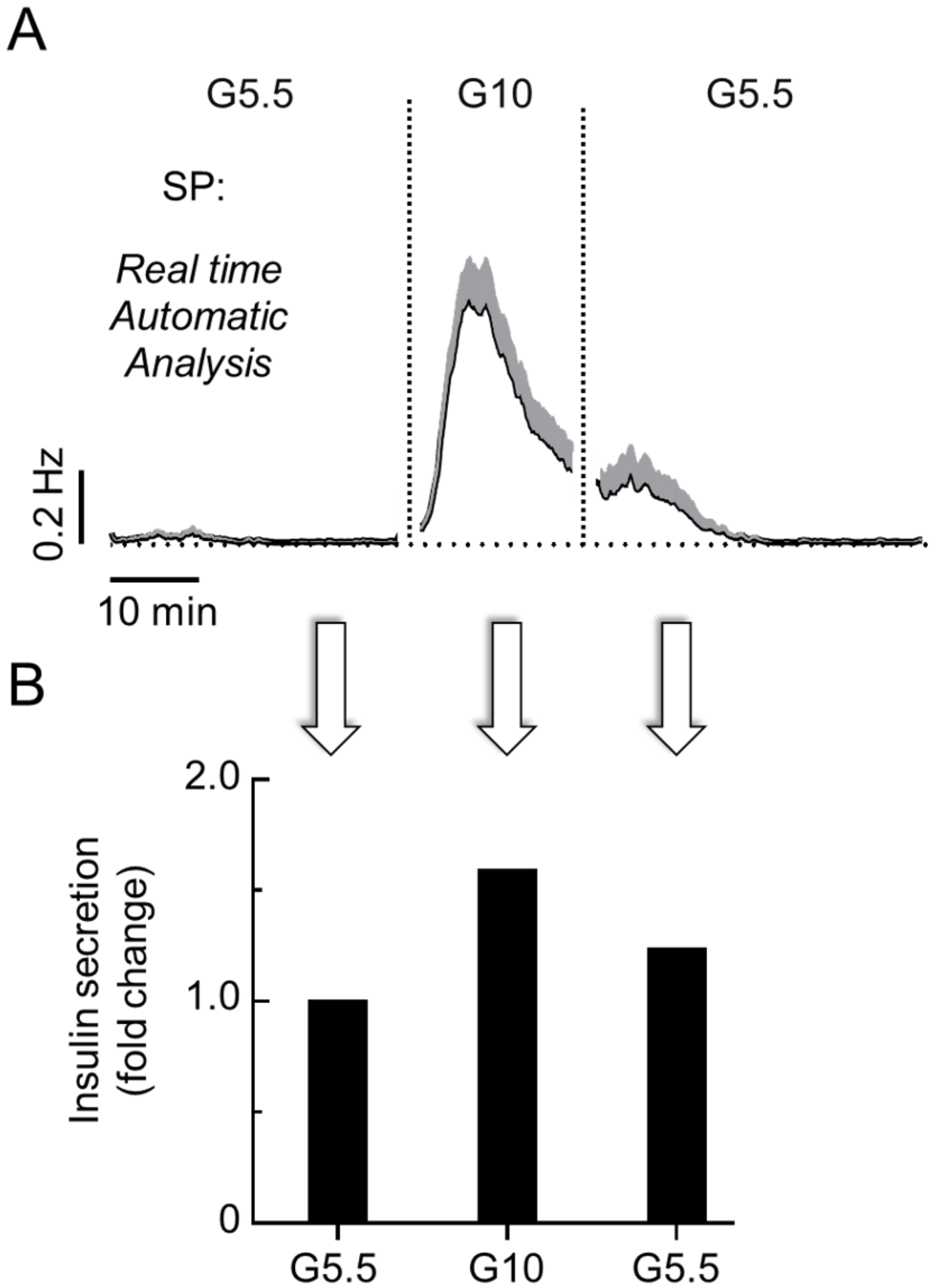


**Supplementary Figure 4**

**Supplementary Figure 4: Combination of automatic microelectronic analysis of SPs and insulin secretion.** (**A**) Real time automatic hardware determination of SP frequencies (means +SEM, n= 29) in BLCs in monolayer when glucose increases from 5.5 to 10 mM and then returns to 5.5 mM. (**B**) Corresponding insulin secretions (relative to the first G5.5).
